# *Coxiella burnetii* blocks intracellular IL-17 signaling in macrophages

**DOI:** 10.1101/367920

**Authors:** Tatiana M. Clemente, Minal Mulye, Anna V. Justis, Srinivas Nallandhighal, Tuan M. Tran, Stacey D. Gilk

## Abstract

*Coxiella burnetii* is an obligate intracellular bacterium and the etiological agent of Q fever. Successful host cell infection requires the *Coxiella* Type IVB Secretion System (T4BSS), which translocates bacterial effector proteins across the vacuole membrane into the host cytoplasm, where they manipulate a variety of cell processes. To identify host cell targets of *Coxiella* T4BSS effector proteins, we determined the transcriptome of murine alveolar macrophages infected with a *Coxiella* T4BSS effector mutant. We identified a set of inflammatory genes that are significantly upregulated in T4BSS mutant-infected cells compared to mock-infected cells or cells infected with wild type (WT) bacteria, suggesting *Coxiella* T4BSS effector proteins downregulate expression of these genes. In addition, the IL-17 signaling pathway was identified as one of the top pathways affected by the bacteria. While previous studies demonstrated that IL-17 plays a protective role against several pathogens, the role of IL-17 during *Coxiella* infection is unknown. We found that IL-17 kills intracellular *Coxiella* in a dose-dependent manner, with the T4BSS mutant exhibiting significantly more sensitivity to IL-17 than WT bacteria. In addition, quantitative PCR confirmed increased expression of IL-17 downstream signaling genes in T4BSS mutant-infected cells compared to WT or mock-infected cells, including the pro-inflammatory cytokines *I11a, Il1b* and *Tnfa*, the chemokines *Cxcl2* and *Ccl5*, and the antimicrobial protein *Lcn2*. We further confirmed that the *Coxiella* T4BSS downregulates macrophage CXCL2/MIP-2 and CCL5/RANTES protein levels following IL-17 stimulation. Together, these data suggest that *Coxiella* downregulates IL-17 signaling in a T4BSS-dependent manner in order to escape the macrophage immune response.

## INTRODUCTION

The intracellular bacterium *Coxiella burnetii* is the etiological agent of Q fever, a zoonotic infectious disease. Initially, Q fever manifests as an acute self-limited flu-like illness. However, patients can develop chronic disease that can be life threatening due to serious clinical manifestations such as endocarditis (1). Furthermore, the current therapy recommended for chronic Q fever requires at least 18 months of doxycycline and hydroxychloroquine treatment (2). An effective vaccine (Q-Vax) has been developed for humans but is currently licensed only in Australia due to adverse effects, especially when administered in previously infected populations (3). In addition, Q fever outbreaks have occurred in several countries, including the Netherlands (4), US (5), Spain (6), Australia (7), Japan (8) and Israel (9), exemplifying how expansive *C. burnetii* infection is worldwide and the need for novel therapeutic targets.

Human infection occurs primarily by inhaling contaminated dust or aerosols, often from close contact with livestock. In the lungs, *C. burnetii* displays tropism for alveolar macrophages, where it forms a phagolysosome-like parasitophorous vacuole (PV) necessary to support bacterial growth (10, 11). *C. burnetii’s* ability to survive and replicate inside the PV, an inhospitable environment for most bacteria, is a unique feature essential for *C. burnetii* pathogenesis. *C. burnetii* exploits the acidic PV pH for metabolic activation (12) and actively manipulates PV fusogenicity and maintenance (13). PV establishment requires translocation of bacterial proteins into the host cell cytoplasm by the *C. burnetii Dot/Icm* (defect in organelle trafficking/intracellular multiplication) type IVB secretion system (T4BSS), closely related to the *Dot/Icm* T4BSS of *Legionella pneumophila* (14). T4BSS effector proteins not only manipulate host vesicular trafficking during PV development, but also other cellular processes such as lipid metabolism, host gene expression, apoptosis, host translation, iron transport, ubiquitination, autophagy and immunity (15, 16). Based on *in silico* prediction, there are more than 100 putative *C. burnetii* T4BSS effector proteins (17–19), but functional data is lacking for the majority of these proteins. In particular, the role of T4BSS effector proteins in manipulating the innate immune response is poorly understood. Recently, the *C. burnetii* T4BSS effector protein IcaA was found to inhibit caspase 11-mediated, non-canonical activation of the nucleotide binding domain and leucine rich repeat containing protein (NLRP3) inflammasome during *C. burnetii* infection (20). Since cytosolic lipopolysaccharide (LPS) is known to activate non-canonical inflammasomes (21, 22), it is possible that *C. burnetti* LPS triggers this pathway, and the bacterium utilizes T4BSS effectors such as IcaA to block this innate immune response. Given the low infectious dose (< 10 organisms) (23), *C. burnetii* certainly inhibits several immediate host cell responses in order to establish infection.

In order to identify new immune response pathways manipulated by *C. burnetii* T4BSS effector proteins, we compared the transcriptome of alveolar macrophages infected with wild type (WT) or a T4BSS mutant *C. burnetii*. We identified a set of inflammatory genes downregulated by *C. burnetii* T4BSS effector proteins, with the IL-17 signaling pathway being one of the top targeted host cell pathways. As IL-17 is a pro-inflammatory cytokine that plays a role in the protective response against a variety of bacterial infections, including the pulmonary intracellular pathogens *Mycoplasma pneumonia, Mycobacterium tuberculosis, Francisella tularensis*, and *Legionella pneumophila* (24–27), we further investigated the role of IL-17 during *C. burnetii* infection. Our data revealed that stimulating the macrophage IL-17 signaling pathway leads to *C. burnetii* killing in a dose-dependent manner, with the T4BSS mutant displaying increased sensitivity compared to WT bacteria. Finally, our findings demonstrated that *C. burnetii* downregulates the IL-17 signaling pathway in macrophages through T4BSS effector proteins.

## RESULTS

### Differentially expressed genes in *C. burnetii*-infected macrophages

In order to identify T4BSS-dependent changes in expression of host genes, we determined the whole transcriptome of murine alveolar macrophages (MH-S) infected with either wild type (WT) *C. burnetii* or a *C. burnetii* mutant lacking *icmD*, an essential component of the T4BSS (14). We previously found minimal differences in PV size and bacterial replication between WT and a T4BSS mutant *C. burnetii* during the first 48 hours of infection of MH-S macrophages (28). Thus, to avoid changes in host cell gene expression that could occur due to PV expansion and bacterial replication after 48 hours, and because *C. burnetii* T4BSS effector protein secretion occurs by four hours post infection (29), we analyzed gene expression at 24 and 48 hours post infection (hpi). By principal components analysis (PCA), global transcription in T4BSS-mutant-infected cells more closely resembled mock-infected cells than WT-infected cells, suggesting that the active T4BSS in WT bacteria drastically alters the host cell response to *C. burnetii* infection (Fig. 1A). The number of differentially expressed genes (DEGs) were determined for each comparison at 24 or 48 hpi, using an absolute fold-change threshold of >1.5 and false discovery rate (FDR) <5% (Supplementary Data Set 1). The largest differences in gene expression at 24 hpi and 48 hpi were for the *icmD* mutant-infected vs. WT-infected comparison (110 DEGs at 24 hpi) and WT-infected vs. mock-infected comparison (116 DEGs at 48 hpi), respectively (Fig. 1B, Supplementary Data Set 1). Unsupervised hierarchical cluster analysis of fold-change values for DEGs (absolute log_2_ fold-change > 0.585; FDR<5%) across all six possible two-way comparisons revealed that the majority of DEGs were upregulated in *icmD* mutant-infected compared to WT-infected cells (*icmD* vs. WT) (Fig. 1C). In contrast, the majority of DEGs were downregulated in WT-infected vs. mock-infected cells (WT vs. mock). Overall there were fewer downregulated genes in the *icmD* mutant-infected cells vs. mock-infected cells (*icmD* vs. mock). This provides evidence that *C. burnetii* T4BSS effector proteins may play a role in the downregulation of host cell genes during the initial stages of infection.

**Figure 1:**
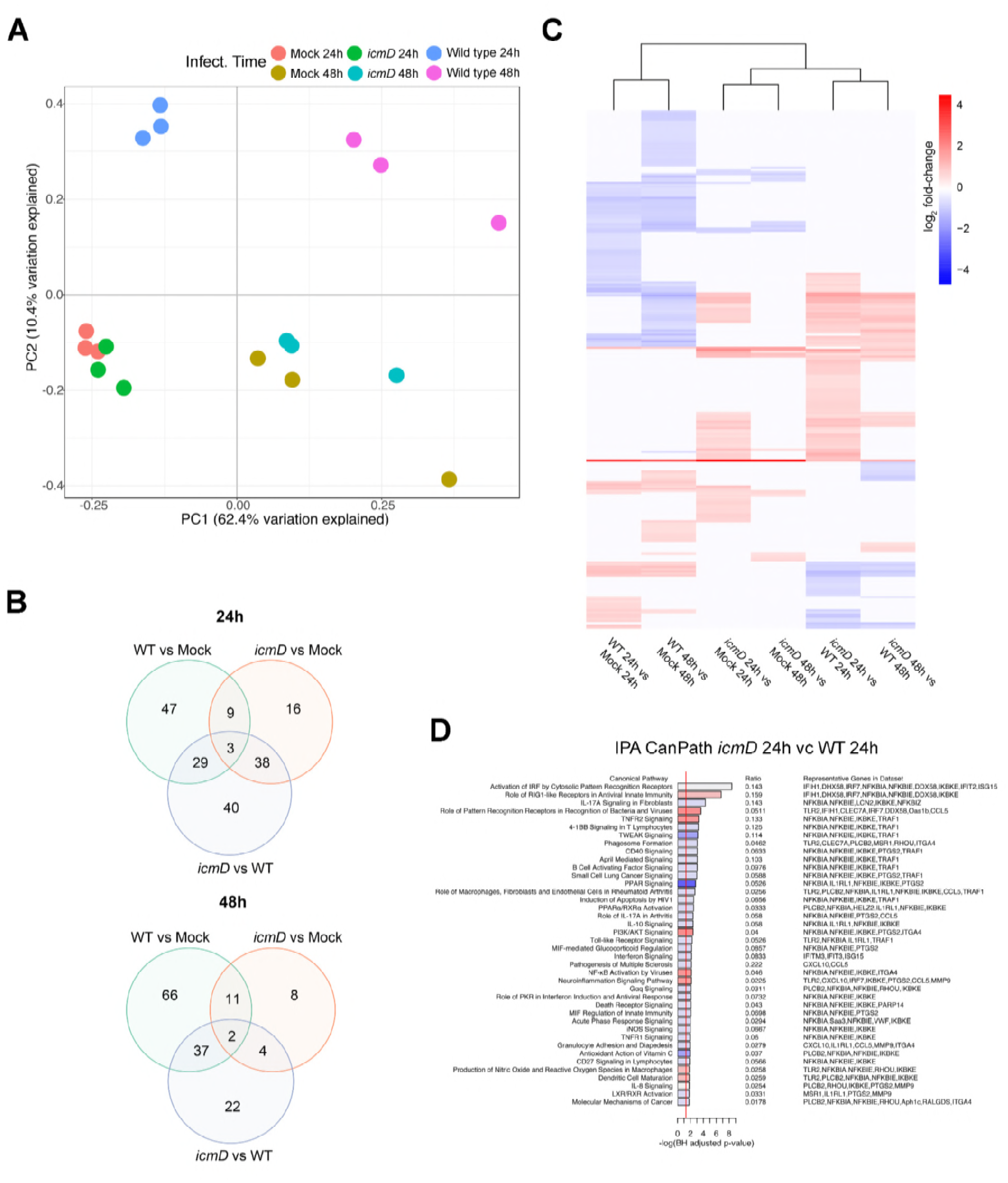
*C. burnetii* infection alters the gene expression profile of alveolar macrophages in a T4BSS-dependent manner. Transcriptome of MH-S macrophages infected with wildtype (WT) or *icmD* T4SS mutant *C. burnetii* at 24 or 48 hpi. **(A)** Principal components analysis (PCA) of genome-wide expression across all RNA-seq samples after normalization of raw data. **(B)** Venn diagram of differentially expressed genes for each comparison for each time point using an absolute log_2_ fold-change (logFC) of >0.585 (1.5-fold in linear space) and a false discovery rate (FDR) <0.05. **(C)** Unsupervised hierarchical clustering heat map of log_2_ fold-change values for all six comparisons using ward. D2 clustering method and euclidean distance. Red intensity indicates increased expression in first group relative to second group, whereas blue intensity indicates decreased expression. **(D)** Ingenuity Pathways Analysis (IPA) using differentially expressed genes for the comparison between WT-infected cells and *icmD*-infected cells at 24 hpi. Red shading indicates increased pathway activity in *icmD* mutant-infected cells, whereas blue shading indicates increased pathway activity in WT-infected cells based on IPA activity z-scores. White shading indicates no activity pattern predicted or could be determined.

To identify biological pathways targeted by *C. burnetii* T4BSS effector proteins, we used two methods: gene set enrichment via CERNO testing (30) using Gene Ontology (GO) annotations as gene sets (Supplementary Fig. 1) and the Ingenuity Pathways Analysis (IPA) using DEGs with an absolute log_2_ fold-change > 0.585 and FDR < 5% as input (Fig. 1D and Supplementary Fig. 2). Both methods revealed differential expression of several immune and inflammatory pathways, including pathogen recognition and activation of Interferon-regulatory factor (IRF) by cytosolic pattern recognition receptors (PRRs) and transmembrane PRRs, signaling pathways induced by the pro-inflammatory cytokines IL-1α and IL-1β, chemokine activity, T cell migration and NF-Kb phosphorylation. In addition, our data indicate that the *C. burnetii* T4BSS significantly downregulates the macrophage type I interferon (IFN) response. This finding supports published data that *C. burnetii* does not induce a robust type I IFN response in macrophages (31). In addition, IL-17 signaling was among the top three overrepresented canonical pathways between mutant and WT infection (Fig. 1D) with upstream regulator analysis predicting activation of IL-17 signaling in *icmD* mutant-infected cells relative to mock-infected cells (Supplementary Fig. 2). Given that IL-17 is known to be an important pro-inflammatory cytokine against several pulmonary pathogens, we specifically tested for differential expression of IL-17 related genes (32) using self-contained gene set testing. The IL-17 gene set was overexpressed in *icmD* mutant-infected macrophages relative to WT-infected macrophages (Supplementary Table 1), suggesting that the *C. burnetii* T4BSS downregulates IL-17 signaling in macrophages.

To validate the transcriptome analysis, we used quantitative reverse transcription PCR (RT-qPCR) of infected macrophages to test expression of IL-17 pathway and other pro-inflammatory genes. RNA was isolated from MH-S macrophages infected with either WT *C. burnetii* or *C. burnetii* mutant lacking *dotA*, another essential component of the T4BSS (33). Like the *icmD* mutant, the *dotA* mutant does not translocate T4BSS effector proteins, allowing us to confirm that gene expression changes are indeed T4BSS-dependent. Lipopolysaccharide (LPS), a potent stimulator of the inflammatory response (34), served as a positive control. Between the WT and *dotA* mutant-infected macrophages, we observed a significant difference in gene expression of the pro-inflammatory genes *Il1a, Il1b* and *Tnfa* (Fig. 2A-C) as well as the IL-17 signaling pathway chemokines *Cxcl2*/*Mip2* and *Ccl5*/*Rantes* (Fig. 2D-E) and the antimicrobial protein Lipocalin-2 (*Lcn2*) (Fig. 2F). These genes were upregulated in the *dotA* mutant-infected macrophages compared to WT-infected macrophages, with more significant differences at 24 hpi compared to 48 hpi. *IL-17A* itself was not differentially regulated in either our RNAseq data or RT-qPCR (data not shown), which is not surprising given that macrophages produce very little IL-17 (35). These data suggest that, during the early stages of macrophage infection, the *C. burnetii* T4BSS may target the IL-17 pathway in order to downregulate expression of several pro-inflammatory genes.

**Figure 2:**
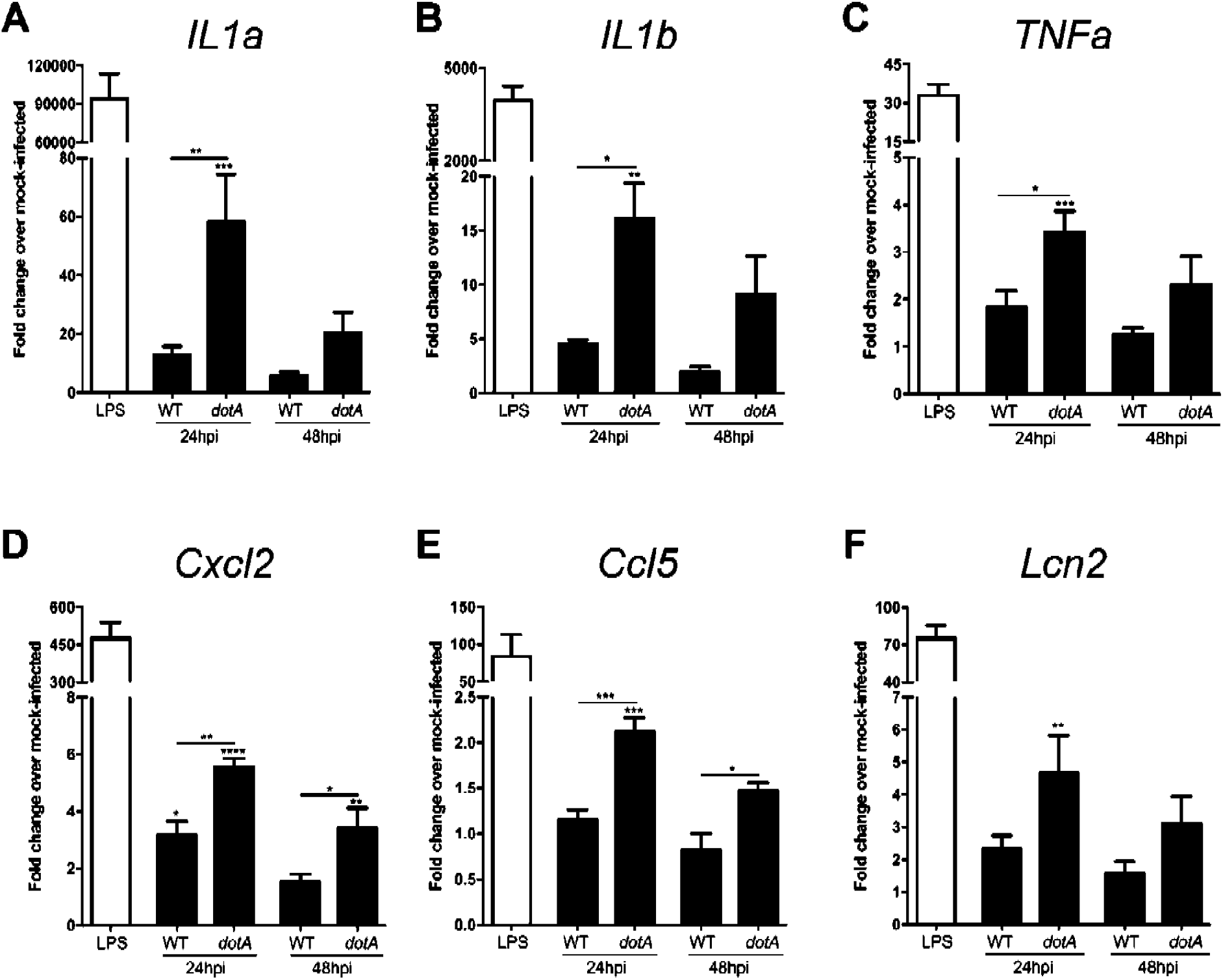
*C. burnetii* T4BSS effector proteins downregulate expression of the IL-17 pathway. Quantitative RT-PCR gene expression analysis of IL-17 pathway genes in macrophages infected with wildtype (WT) or *dotA* mutant *C. burnetii* for 24 or 48 hpi. Mock-infected macrophages treated with LPS (100 ng/ml) for 24 hours were used as a positive control. Individual genes were normalized to *Gapdh* and the fold change in expression over mock-infected cells determined. When compared to WT-infected cells, macrophages infected with *dotA* mutant *C. burnetii* have higher expression of the IL-17 pathway genes **(A)***IL1a*, **(B)** *IL1b*, **(C)***Tnfa*, **(D)***Cxcl2*, **(E)***Ccl5*, and **(F)***Lcn2*. Error bars show the average +/- SEM of three independent experiments, performed in biological duplicate with three technical replicates. *p<0.05, **p<0.01, ***p<0.005 and ****p<0.001 as determined by one-way ANOVA with Tukey’s posthoc test compared to mock-infected or between WT and *dotA* mutant-infected cells.

### *C. burnetii* downregulates CXCL2/MIP-2 and CCL5/RANTES expression in a T4BSS-dependent manner

A number of studies have shown that IL-17 plays an important role in the innate immune response against bacteria by stimulating secretion of multiple chemokines. Within the context of infection, these chemokines recruit macrophages, neutrophils, and lymphocytes to the infection site, thereby enhancing inflammation. To validate the gene expression changes between macrophages infected with WT or the T4BSS mutant, we measured secretion of CXCL2/MIP-2 and CCL5/RANTES at 24 or 48 hpi using ELISA. We observed a significant difference in CXCL2/MIP-2 and CCL5/RANTES protein levels between the WT and *dotA* mutant-infected macrophages, with a 3-fold increase of both cytokines in the *dotA* mutant-infected macrophages (Fig. 3A-B), confirming the gene expression data. While CXCL2/MIP-2 was significantly higher at both 24 and 48 hpi, we only detected a difference in CCL5/RANTES expression at 24 hpi (Fig. 3B).

**Figure 3:**
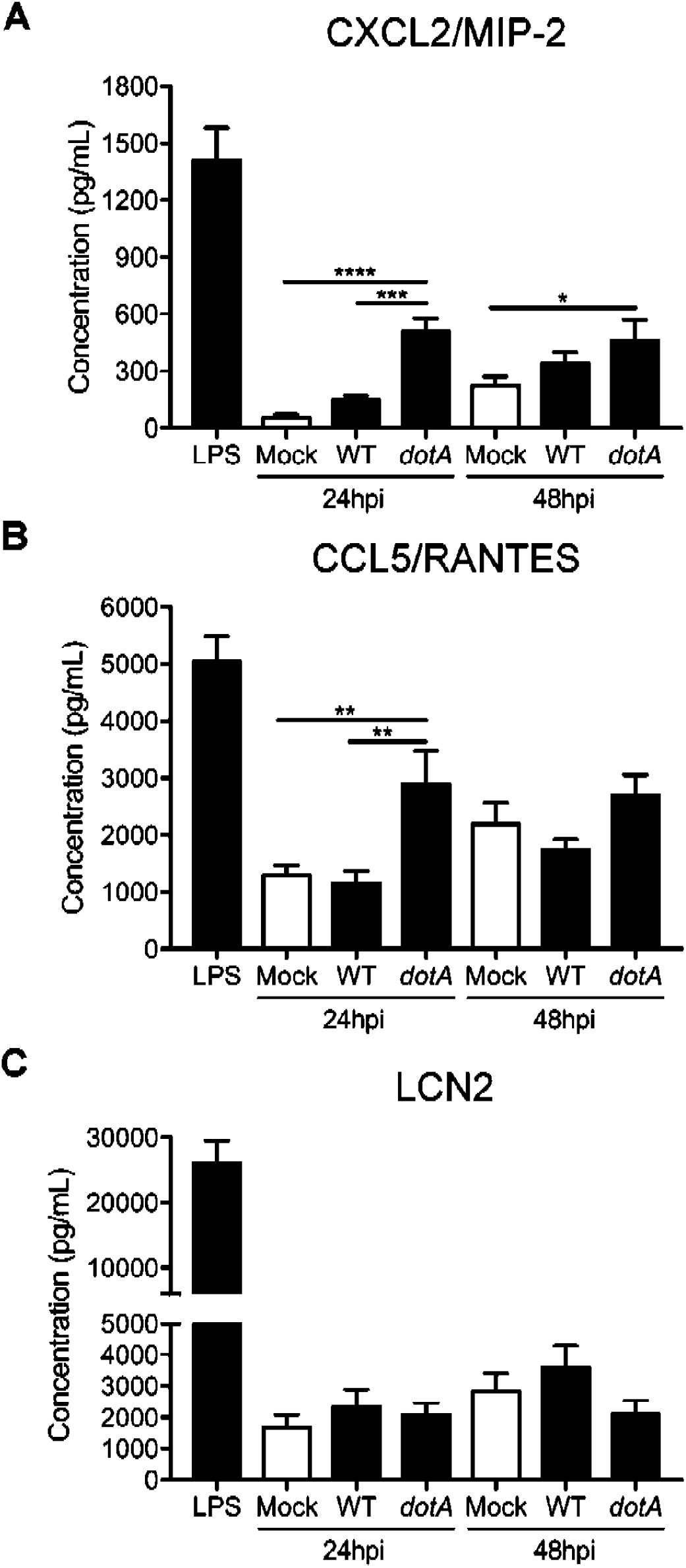
The *C. burnetii* T4BSS decreases secretion of CXCL2/MIP-2 and CCL5/RANTES in infected MH-S macrophages. ELISA protein quantitation of CXCL2/MIP-2, CCL5/RANTES and Lipocalin-2 (LCN2) in the supernatant of macrophages infected with WT or *dotA* mutant *C. burnetii* at 24 or 48 hpi. Cells treated with LPS (100ng/ml) were used as a positive control. Compared to mock-infected and WT-infected macrophages at 24 hpi, *dotA* mutant-infected macrophages have increased secretion of **(A)** CXCL2/MIP-2 and **(B)** CCL5/RANTES, but not **(C)** LCN2. Shown are the means +/- SEM from three independent experiments done in duplicate. *p<0.05, **p<0.01, ***p<0.005 and ****p<0.001 as determined by one-way ANOVA with Tukey’s posthoc test.

LCN2 expression is strongly induced by IL-17 and blocks catecholate-type siderophores of gram-negative bacteria, preventing the bacteria from scavenging free iron required for bacterial growth (36, 37). While *Lcn2* gene expression was differentially regulated (Fig. 2F), we did not observe a significant difference in secreted LCN2 protein between the WT versus *dotA* mutant-infected macrophages at either 24 or 48 hpi (Fig. 3C). These conflicting data may be due to post-translational regulation of LCN2 (38). However, our data does suggest that the *C. burnetii* T4BSS downregulates macrophage secretion of the chemokines CXCL2/MIP-2 and CCL5/RANTES during infection.

### *C. burnetii* T4BSS effector proteins impair IL-17-stimulated CXCL2/MIP-2 and CCL5/RANTES secretion

The chemo-attractant CXCL2/MIP-2 is typically secreted by monocytes and macrophages and recruits neutrophils required for pathogen clearance (39). Furthermore, CCL5/RANTES, which is secreted by lymphocytes, macrophages, and endothelial cells, also recruits and activates leukocytes (40). We first confirmed that IL-17 upregulates CXCL2/MIP-2 and CCL5/RANTES in MH-S alveolar macrophages by treating uninfected macrophages with recombinant mouse IL-17A and analyzing the cell-free supernatant by ELISA. In uninfected macrophages, CXCL2/MIP-2 and CCL5/RANTES increased 14-fold and 2-fold, respectively, following IL-17A treatment for 24 hours (Fig. 4A-B). To test if *C. burnetii* T4BSS effector proteins block IL-17-stimulated chemokine secretion, WT or *dotA* mutant-infected macrophages were treated with IL-17A for 24 hours. In IL-17 stimulated macrophages infected with WT bacteria, CXCL2/MIP-2 decreased 2.2-fold compared to stimulated mock-infected macrophages, while CCL5/RANTES decreased 1.6-fold (Fig. 4A-B), suggesting that WT *C*. *burnetii* blocks IL-17-induced chemokine secretion. Further, *dotA* mutant *C. burnetii* did not block IL-17A-stimulated CXCL2/MIP-2 and CCL5/RANTES (Fig. 4A-B). These data suggest that the *C. burnetii* T4BSS impairs IL-17 signaling in macrophages, including secretion of CXCL2/MIP-2 and CCL5/RANTES.

**Figure 4:**
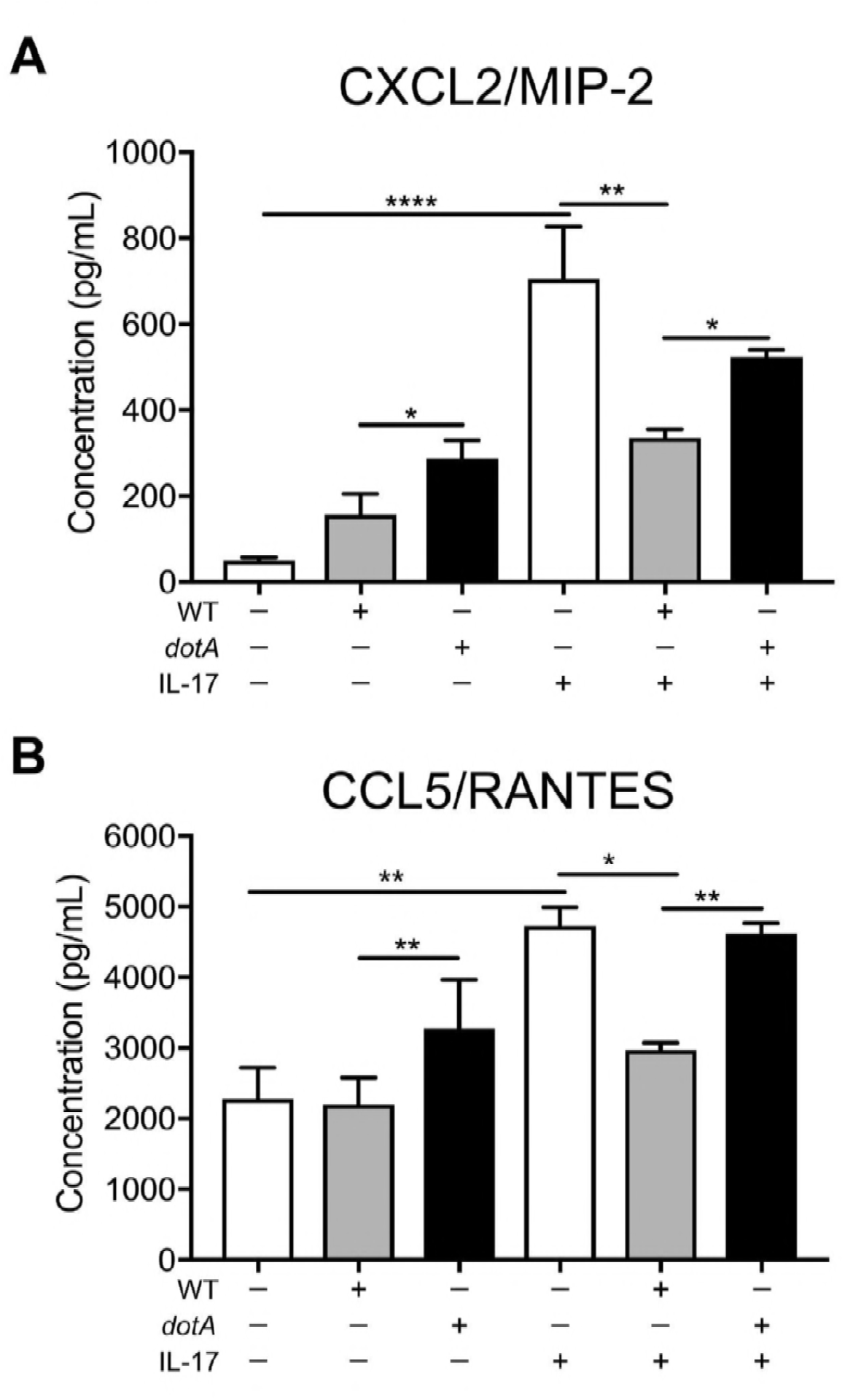
*C. burnetii* T4BSS blocks IL-17 stimulated secretion of CXCL2/MIP-2 and CCL5/RANTES. ELISA quantitation of **(A)** CXCL2/MIP-2 and **(B)** CCL5/RANTES protein levels after IL-17A (100 ng/ml) treatment of mock-infected, WT-infected, or *dotA* mutant-infected macrophages at 24 hpi. The means +/- SEM of three individual experiments, performed in duplicate are shown. *p<0.05, **p<0.01, and ****p<0.001 as determined by one-way ANOVA with Tukey’s posthoc test compared to mock-infected, or between WT and *dotA* mutant-infected cells.

### Triggering the macrophage IL-17 pathway is bactericidal

Several studies have demonstrated that IL-17 plays a protective role for the host during bacterial infections (32, 41–43). To evaluate if IL-17 affects *C. burnetii* viability in macrophages, we treated infected macrophages at 24 or 48 hpi with recombinant mouse IL-17A and enumerated viable bacteria 24 hours later using a fluorescent infectious focus-forming unit (FFU) assay in Vero cells (44). IL-17 stimulation decreased *C. burnetii* viability at 24 and 48 hpi in a dose dependent-manner, with a ∼40% decrease at the highest concentration (Supplementary Fig. 3). IL-17A treatment did not affect the macrophage viability (data no shown). Interestingly, *C. burnetti* appears more resistant to IL-17 stimulation at 48 hpi compared to 24 hpi, as low concentrations of IL-17 (50 ng/ml) led to significant loss of the bacteria viability only at 24 hpi (Supplementary Fig. 3A-B). To determine if the T4BSS is related to bacteria susceptibility to IL-17, we infected macrophages with either WT or *dotA C. burnetii*, stimulated with IL-17A, and measured bacteria viability by colony-forming unit (CFU) assay on agarose plates (45). We observed a stronger bactericidal effect of IL-17 on *dotA* mutant *C. burnetii* compared to WT *C. burnetii*, as the presence of 25 and 12.5 ng/ml of IL-17 led to 47% and 39% loss of *dotA* mutant *C. burnetii* viability, respectively, but did not affect WT *C. burnetii* viability (Fig. 5A and Supplementary Fig. 4). To further assess the specificity of IL-17 activity, we treated infected cells with IL-17 (50 or 100 ng/ml) in the presence or absence of an antibody that blocks the IL-17 receptor. The IL-17 bactericidal effect was significantly neutralized by blocking the IL-17 receptor, as the co-treatment with IL-17 and the anti-IL-17 receptor antibody rescued over 30% of the bacteria viability when compared to IL-17 treated infected cells (Fig. 5B and Supplementary Fig. 4).

In order to validate the bactericidal effect of IL-17 in primary cells, human monocyte-derived macrophages (hMDMs) were infected with WT or *dotA C. burnetii*, treated at 24 hpi with recombinant human IL-17A, and bacterial viability was measured after 24 hours by CFU assay. Confirming our results obtained in MH-S cells, the *dotA* T4BSS mutant viability decreased with IL-17 treatment (Fig. 5C and Supplementary Fig. 4), with a 50% decrease in viable bacteria at 100 ng/ml. However, IL-17 had no effect on WT *C. burnetii* in primary hMDMs. Together, these data suggest that activation of the IL-17 signaling pathway in macrophages kills intracellular *C. burnetii*, with the *C*. *burnetii* T4BSS playing a protective role.

**Figure 5:**
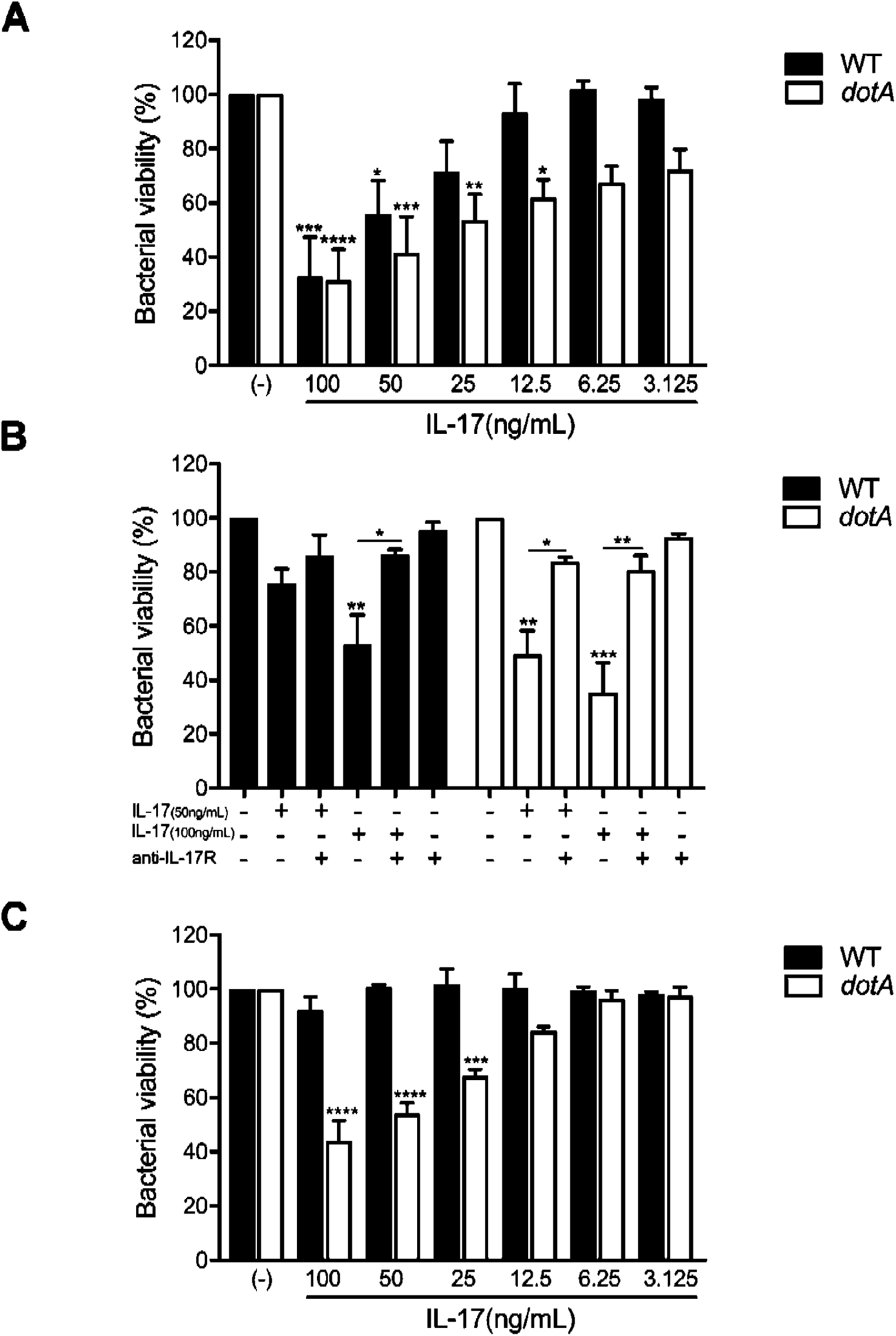
Activating the macrophage IL-17 pathway can kill *C burnetii*, but the T4BSS effector proteins play a protective role. MH-S cells were infected for 24 hours, followed by treatment for 24 hours with either **(A)** IL-17 alone or **(B)** IL-17 and an IL-17A receptor blocking antibody (2 μg/ml). hMDMs **(C)** were infected for 24 hours and treated with human IL-17A for 24 hours. Viable bacteria were quantitated using an agarose-based colony forming unit (CFU) assay, and the loss of bacterial viability calculated by dividing the number of WT or T4SS mutant bacteria in treated samples to their respective untreated (-) samples. Compared to WT bacteria, the *dotA* T4BSS mutant is more sensitive to IL-17 in both MH-S and hMDMs, and viability of both WT and *dotA* mutant *C. burnetii* can be recovered by blocking IL-17 receptor signaling. Error bars indicate the mean +/- SEM from four individual experiments. *p<0.05, **p<0.01, ***p<0,005 and ****p<0.001 as determined by one-way ANOVA with Dunnett’s posthoc test compared to untreated controls.

## DISCUSSION

The innate immune response relies on pathogen detection by pattern recognition receptors which activate signaling pathways and trigger an inflammatory response (46). While essential to protect the host, pathogens such as *C. burnetii* have evolved strategies to overcome the host innate immune response (47, 48). Despite being sequestered in a growth-permissive vacuole, *C. burnetii*

T4BSS effector proteins manipulate a variety of host cell signaling processes, including the innate immune responses of inflammasome-mediated pyroptotic and apoptotic cell death (20, 49–51). To identify potential targets of *C. burnetii* T4BSS effector proteins, we compared the transcriptome of murine alveolar macrophages infected either with WT or T4BSS mutant *C. burnetii*. We identified several inflammatory pathways downregulated by *C. burnetii* T4BSS effector proteins, including IL-17 signaling. Previous studies demonstrated that IL-17 plays a protective role against several pathogens, including *L. pneumophila*, the closest pathogenic relative to *C. burnetii*(25–27, 41, 42). We found that *C. burnetti* downregulates the macrophage IL-17 signaling pathway in a T4BSS-dependent manner, protecting the bacteria from IL-17-mediated killing by the macrophage and blocking secretion of pro-inflammatory chemokines. To our knowledge, this is the first demonstration of a pathogenic bacteria directly downregulating intracellular macrophage IL-17 signaling.

Previous studies demonstrated that *C. burnetii* infection leads to secretion of the pro-inflammatory cytokines TNFα and IFNγ, with both cytokines playing critical roles in restricting *C. burnetii* replication (52–54). In our studies, gene expression analysis during the early stages of infection revealed striking differences in the immunological response to WT and T4BSS mutant *C. burnetii*, with *C. burnetii* T4BSS mutant-infected macrophages having a stronger pro-inflammatory response. For example, the pro-inflammatory genes *Il1a, Il1b* and *Tnfa* are expressed at higher levels in macrophages infected with T4BSS mutant *C. burnetii*, compared to WT-infected macrophages. Bacterial-driven downregulation of these and other pro-inflammatory cytokines would benefit the bacteria in establishing infection. In support of our data, *C. burnetii* infection of primary macrophages does not activate caspase-1 (20), an enzyme required for the production of the pro-inflammatory cytokines IL-1β and IL-1α (55, 56). Interestingly, *C. burnetii* does not directly inhibit caspase-1 activation but appears to interfere with upstream signaling events, including blocking TNFα signaling (20, 57). However, a recent study did not detect significant differences in TNFα production in murine bone-marrow derived macrophages infected with WT *C. burnetii* or *icmL C. burnetii*, a mutant with non-functional T4BSS (31). These apparently conflicting data may be explained by the use of C57BL/6 mice in the latter study; C57BL/6 mice, in contrast to other inbred mouse strains, are not permissive for intracellular *C. burnetii* replication due to the large amount of TNFα produced upon toll-like receptor (TLR) stimulation (31, 58–60). Further experimentation is required to elucidate the mechanism(s) behind *C. burnetii* T4BSS-mediated downregulation of macrophage pro-inflammatory response.

Pathogen-associated molecular patterns (PAMPs) are sensed by different PRRs, which activate IRFs and initiate key inflammatory responses including transcription of type I interferons (IFN) and IFN-inducible genes (61, 62). Type I IFN can be induced by many intracellular bacterial pathogens, either via recognition of bacterial surface molecules such as LPS, or through stimulatory ligands released by the bacteria via specialized bacterial secretion systems (63). Our transcriptome analysis revealed *C. burnetii* T4BSS-mediated downregulation of macrophage IRF activation by cytosolic and transmembrane PRRs. A recent study found that *C. burnetii* does not trigger cytosolic PRRs or induce robust type I IFN production in mouse macrophages (31). Additionally, IFN-α receptor-deficient (IFNAR^-/-^) mice were protected from *C. burnetii* infection, suggesting that type I IFNs are not required to restrict bacterial replication (31, 64). However, delivery of recombinant IFN-α to the lung of *C. burnetii*-infected mice protected against bacterial replication, revealing a potential role of type I IFN in control of *C. burnetii* infection in the lung (64). Interestingly, type I IFN is induced during *L. pneumophila* infection and plays a key role in macrophage defense by restricting intracellular bacterial replication (65, 66). However, to counteract this host immune response, the *L. pneumophila* T4SS effector protein SdhA suppresses induction of IFN through an unknown mechanism (67). Similarly, our data suggests that *C. burnetii* T4BSS effector proteins negatively modulate the type I IFN response in alveolar macrophages, most likely as a bacterial immune evasion mechanism.

In addition to pro-inflammatory cytokines and PRRs, we discovered an important role for the cytokine IL-17 during *C. burnetii* infection of macrophages. The protective role of IL-17 against extracellular bacteria has been extensively studied; additionally, IL-17 can be critical for the full immune response leading to the control of intracellular bacteria (32, 42, 43, 68). IL-17 is produced by T helper 17 (Th17) cells, γδ T cells and invariant natural killer T (iNKT) cells (69). In the lung, γδ T cells have been implicated as a primary source of early IL-17 production in several *in vivo* models of infection (70), which may have implications for *C. burnetii* lung infection. Exogenous IL-17 binds the IL-17 receptor on the surface of the macrophage, triggering chemokine secretion, neutrophil recruitment, and a Th1 response, thus enhancing bacterial clearance (26, 27, 71, 72). By both gene expression and protein analysis, we found that *C. burnetii* downregulates IL-17-stimulated chemokine secretion in macrophages in a T4BSS-dependent manner. A previous study found that following *C. burnetii* aerosol infection in mice, neutrophils are not present in the airways until 7 days post infection, though the mechanism of this delay remains unknown (73). Further, neutrophils play a critical role in inflammation and bacterial clearance following intranasal *C. burnetii* infection, but it is unknown whether neutrophils directly kill the bacteria or serve to enhance the immune response (74). Based on our findings in alveolar macrophages, we hypothesize that *C. burnetii* T4BSS effector proteins downregulate the IL-17 pathway to suppress chemokine secretion as a mechanism to avoid neutrophil recruitment at early stages of infection. This could be an important immune evasion strategy that enables the bacteria to establish long-term persistence. In addition to chemokines, the IL-17-stimulated protein LCN2 may also be downregulated by *C. burnetii*. LCN2 is a siderophore-binding antimicrobial protein that can limit bacterial growth by iron restriction.

A previous study demonstrated that *C. burnetii*-infected IL-17 receptor knockout mice had a similar bacterial burden in the spleen and lung as infected WT mice, suggesting that IL-17 does not play an essential role during *C. burnetii* infection (74). In contrast, our *in vitro* studies revealed that activating the IL-17 signaling pathway in macrophages can directly kill intracellular *C. burnetii*. Further, the *C. burnetii* T4BSS appears to play a protective role, presumably by blocking the intracellular signaling pathway triggered by IL-17 binding to the IL-17R. Our data may explain the lack of phenotypic changes in IL-17 receptor knockout mice infected by WT *C. burnetii*, as the intracellular signaling pathway is not activated in the absence of the IL-17 receptor.

During acute *C. burnetii* infection in humans, the number of γδ T cells rise significantly in the peripheral blood of patients (75). Given that γδ T cells can secrete large amounts of IL-17 (76), it is possible that the downregulation of the intracellular IL-17 signaling by T4BSS effector proteins might be an essential mechanism of immune evasion that allows *C. burnetii* persistence. IL-17 activates common downstream pathways in macrophages, including NF-κB (Nuclear factor-κB) and MAPKs (mitogen-activated protein kinases) (77, 78). Our transcriptome data suggests that the *C. burnetii* T4BSS downregulates the IL-17 canonical NF-κB signaling pathway, including *Il17ra, Il17rc, Traf6, Nfkb1* and *Nfkb2*. This hypothesis is consistent with a recent study that found that *C. burnetii* can modulate NF-κB canonical pathway through the T4BSS (79). NF-κB activation correlates with enhanced expression of inducible nitric oxide synthase (iNOS) (80) and NADPH oxidase (NOX) (81), which generate nitric oxide (NO) and reactive oxygen species (ROS), respectively. Both NO and ROS are signature molecules for M1 macrophages (82), while *C. burnetii*-infected macrophages exhibit a M2-polarization that is unable to control bacterial replication (83). As IL-17 alters macrophage polarization (84), one potential mechanism is that IL-17 polarizes toward M1 phenotype, triggering ROS and NO leading to *C. burnetii* killing. In addition, as IFNγ plays a clear role in *C. burnetii* killing (53, 54), the IL-17-bactericidal effect might be related to IFNγ, as IL-17 can induce an IFNγ response (85). Further experimentation is needed to not only identify the *C. burnetii* T4BSS effector protein modulating IL-17 signaling in macrophages, but also how IL-17 leads to *C. burnetii* death inside of macrophages.

In summary, this study suggests that *C. burnetii* employs the T4BSS to downregulate IL-17 signaling in macrophages during the early stages of infection. This has important implications in both controlling the pro-inflammatory response elicited by the macrophages, as well as avoiding direct killing by the macrophage. Further studies identifying the bacterial T4BSS effector proteins involved in this mechanism and elucidating how IL-17 kills *C. burnetii* will give new insight into immune evasion by *C. burnetii*.

## MATERIALS AND METHODS

### Bacteria and mammalian cells

*Coxiella burnetii* Nine Mile Phase II (NMII, clone 4, RSA439) were purified from Vero cells (African green monkey kidney epithelial cells, CCL-81; American Type Culture Collection, Manassas, VA) and stored as previously described (86). For all experiments *C. burnetii* NMII wild type (WT), *icmD* (14) and *dotA*(33) mutants were grown for 4 days in ACCM-2, at 37°C in 2.5% O_2_ and 5% CO_2_, washed twice with phosphate buffered saline (PBS) and stored as previously described (87). Murine alveolar macrophages (MH-S; CRL-2019 ATCC) were maintained in growth media consisting of RPMI (Roswell Park Memorial Institute) 1640 medium (Corning, New York, NY, USA) containing 10% fetal bovine serum (FBS, Atlanta Biologicals, Norcross, GA, USA) at 37°C and 5% CO_2_. The multiplicity of infection (MOI) for each bacteria stock was optimized for each bacteria stock and culture vessel for a final infection of approximately 1 internalized bacterium per cell. To obtain human monocyte derived macrophages (hMDM), peripheral blood mononuclear cells were isolated from buffy coats (Indiana Blood Center) using Ficoll-Paque (GE Healthcare # 17144002). Monocytes were isolated from lymphocytes by positive selection using CD14 magnetic beads (Dynabeads^®^ FlowComp™ Human CD14 – catalog # 11367D). Following isolation, monocytes were cultured for seven days with RPMI 1640 medium containing 10% FBS, 100 mg/ml penicillin/streptomycin and 50 ng/ml human macrophage colony-stimulating factor (M-CSF; ThermoFisher Scientific, catalog # 14–8789-62). 24 hours prior infection, the media containing antibiotics and M-CSF was replaced with RPMI 1640 containing 10% FBS.

### RNA sequencing

MH-S cells (4×10^5^ cells per well of a 6-well plate) were mock-infected or infected with WT or *icmD* mutant *C. burnetii*, with three replicates per condition. Total RNA was isolated at 24 and 48 hpi using RNeasy Plus Mini Kit (Qiagen). RNA samples had an RNA integrity number >7, as determined on an Agilent 2100 Bioanalyzer. RNA-seq libraries were prepared using the ScriptSeq Complete kit (Illumina, Inc) according to the manufacturer’s instructions. Libraries were sequenced at 30 million reads per sample on an Illumina NextSeq platform with read lengths of 75 bp by Indiana University Bloomington Center for Genomics and Bioinformatics and mapped to the mouse reference genome mm10 by the Indiana University Center for Computation Biology and Bioinformatics. RNA processing and sequencing were performed as a single batch. The median library size (mapped reads) was 17.8 million reads with a minimum of 13.4 million reads.

### Gene expression analysis

RNAseq differential gene expression (DGE) analysis was performed using the edgeR package (version 3.16.5) in R (version 3.3.3). After filtering genes with low expression across a majority of samples, trimmed mean of M values (TMM) normalization was applied to the remaining 9400 genes. Expression data for these genes were converted to log counts-per-million (logCPM) for data visualization with principal components analysis (PCA) plots. DGE analysis was performed using the glmLRT function as 2-way comparisons between the three classes using the following model matrix formula: ∼0+Infection_Time where Infection_Time is combined factor variable consisting of cell treatment and time point (six levels). The fold change in gene expression was determined by comparing wild type or *icmD* mutant-infected to mock-infected cells or each other at either 24h or 48h (six different comparisons). Differential expression of functional pathways was assessed by two methods using the list of differentially expressed genes (DEGs) for each comparison: 1) gene enrichment analysis using CERNO testing in the *tmod* package (version 0.31) (30) in R with Gene Ontology (GO, C5 in MSigDB (88)) annotations as gene sets and all DEGs without cut-off criteria but ranked by ascending P values and 2) Ingenuity Pathways Analysis (version 42012434) using DEGs with |log_2_FC|>0.585 (1.5 in linear space) and FDR<5% as input. In addition, self-contained gene set testing for enrichment of IL-17 related genes (32) was also performed using the roast. DGElist function in the *edgeR* package and the following gene list: *Ccl5, Il17rc, Lcn2, Traf6, Il17ra, Nfkb1, Nfkb2, Ccl2*, and *Ccl3*.

### Quantitative gene expression by real time-PCR (αRT-PCR)

MH-S cells (2×10^5^ cells per well of a 6-well plate) were mock-infected or infected with WT or *dotA* mutant *C. burnetii* in 0.5 ml growth media for 2 hours at 37°C and 5% CO_2_, washed extensively with PBS and incubated in 2 ml of growth media. Cells treated with LPS (100 ng/ml) from *Escherichia coli* O111:B4 (Sigma, catalog # L4392) were used as a positive control. RNA was isolated using the RNeasy Plus Mini Kit at 24 and 48 hpi, analyzed for quantity and A260/280 ratio (Implen NanoPhotometer), and cDNA generated using Super Script III First-strand synthesis system kit (Invitrogen). Real time PCR using Luminaris™ Color HiGreen qPCR Master Mix (ThermoScientific) was done on a Bio-Rad CFX Connect Real-Time System according to manufacturer’s instructions. Mouse specific primers were (5′ to 3′): *Il1b*, forward: TGTAATGAAAGACGGCACACC; reverse: TCTTCTTTGGGTATTGCTTGG; *Il1a*, forward: CGCTTGAGTCGGCAAAGAAAT; reverse: ACAAACTGATCTGTGCAAGTCTC; *Tnfa*, forward: TTCTGTCTACTGAACTTCGGG; reverse: GTATGAGATAGCAAATCGGCT; CCL5/RANTES, forward: ACTCCCTGCTGCTTTGCCTAC; reverse: ACTTGCTGGTGTAGAAATACT; CXCL2/MIP-2, forward: CGCTGTCAATGCCTGAAGAC; reverse: ACACTCAAGCTCTGGATGTTCTTG; *Lcn2*, forward: TTTCACCCGCTTTGCCAAGT; reverse: GTCTCTGCGCATCCCAGTCA; GAPDH, forward: AAGGTCATCCCAGAGCTGAA; reverse: CTGCTTCACCACCTTCTTGA. The relative levels of transcripts were calculated with the ΔΔCt method using *Gapdh* as the internal control. The relative levels of mRNA from the mock-infected samples were adjusted to 1 and served as the basal control value. Each experiment was done in biological duplicate, and qPCR performed on three separate cDNA preparations from each RNA.

### ELISA

CXCL2-MIP-2, CCL5-RANTES and Lipocalin-2 protein levels in cell-free supernatants were measured by ELISA (R&D Systems, Minneapolis, MN) according to the manufacturer’s instructions. In brief, MH-S cells (5×10^4^ cells per well of a 24-well plate) were plated and allowed to adhere overnight. The cells were then mock-infected or infected with WT or *dotA* mutant *C. burnetii* in 0.25 ml growth media for 2 h at 37°C and 5% CO_2_, washed extensively with PBS and incubated in 0.5 ml growth media. To examine the IL-17 pathway expression in infected cells, the cells were pre-treated with 100 ng/ml of IL-17A recombinant mouse protein (ThermoFisher, catalog # PMC0174) for 24 hours, and then infected as previously described. LPS-treated cells (100 ng/ml; Sigma catalog number L4391) were used as a positive control. The cell supernatant was collected at 24 or 48 hpi, centrifuged at 20,000×g for 10 min, and analyzed by ELISA. Each experiment was performed in biological duplicate with two technical replicates.

### *C. burnetii* viability by fluorescent infectious focus-forming unit (FFU) and colony-forming unit (CFU) assays

MH-S cells (5×10^4^ cells per well of a 24-well plate) were plated and allowed to adhere overnight, while monocytes (1×10^5^cells per well of a 24 well plate) were plated and differentiated to hMDMs for seven days. The cells were then mock-infected or infected with WT or *dotA* mutant *C. burnetii* in 0.25 ml growth media for 2 h at 37°C, 5% CO_2_, washed extensively with PBS and incubated in 0.5 ml growth media. At the indicated time points, the cells were treated with different concentrations of either recombinant mouse IL-17A or recombinant human IL-17A (ThermoFisher, catalog # 14–8179-62) for 24 hours. The cells were lysed in sterile water for 5 min and analyzed by FFU assay as previously described (89). For the CFU assay, the released bacteria were diluted 1:5 in ACCM-2 and plated in 2-fold serial dilutions onto 0.25% ACCM-2 agarose plates (45). Plates were incubated for 7–9 days at 37°C, 2.5% O_2_ and 5% CO_2_ and the number of colonies counted to measure bacteria viability. Each experiment was performed in biological duplicate and spotted in triplicate.

### Antibody neutralization

MH-S cells (5×10^4^) were mock-infected or infected with WT or *dotA* mutant *C. burnetii*, in a 24-well plate. At 24 hpi, the cells were treated with 50 or 100 ng/ml of IL-17A, in the presence or absence of 2 μg/ml of anti-IL-17Ra monoclonal antibody (ThermoFisher Scientific, catalog # MAB4481), for 24 hours. Bacteria were released by water lysis and analyzed by CFU assay as described above. Each experiment was performed in biological duplicate and spotted in triplicate.

### Data analysis

Statistical analyses were performed using ordinary one-way ANOVA with Dunnett’s or Turkey’s multiple comparisons tests in Prism 7 (GraphPad Software, Inc La Jolla, CA).

## ACKLOWLEDGEMENTS

We thank Mark Kaplan and Dhritiman Samanta for helpful discussions, and James Ford, Hongyu Gao and Yunlong Liu for assistance with RNA sequencing and bioinformatics. This research was supported by the National Institute of Allergy and Infectious Diseases, NIH (AI121786 to SDG; 5K08AI125682 to TMT).

